# Multimodal single-cell profiling reveals neuronal vulnerability and pathological cell states in focal cortical dysplasia

**DOI:** 10.1101/2024.05.29.596419

**Authors:** Isabella C. Galvão, Manuela Lemoine, Lauana A. Messias, Patrícia A.O.R.A. Araújo, Jaqueline C. Geraldis, Clarissa L. Yasuda, Marina K. M. Alvim, Enrico Ghizoni, Helder Tedeschi, Fernando Cendes, Fabio Rogerio, Iscia Lopes-Cendes, Diogo F. T. Veiga

**Affiliations:** Department of Translational Medicine, School of Medical Sciences, University of Campinas, Campinas (UNICAMP), Brazil; The Brazilian Institute of Neuroscience and Neurotechnology (BRAINN), Campinas, Brazil; Department of Neurology, School of Medical Sciences, University of Campinas (UNICAMP), Campinas, SP, Brazil; Department of Pathology, School of Medical Sciences, University of Campinas (UNICAMP), Campinas, Brazil

**Keywords:** focal cortical dysplasia, cortical malformations, single-cell sequencing, multi-omics, gene expression, chromatin accessibility, microglia, reactive astrocytes

## Abstract

Focal Cortical Dysplasia (FCD) is a neurodevelopmental condition characterized by malformations of the cerebral cortex that often cause drug-resistant epilepsy. In this study, we performed multi-omics single-cell profiling to map the chromatin accessibility and transcriptome landscapes of FCD type II, generating a comprehensive multimodal single-cell dataset comprising 61,525 cells from 11 clinical samples of lesions and controls. Our findings revealed profound chromatin, transcriptomic, and cellular alterations affecting neuronal and glial cells in FCD lesions, including the selective loss of upper-layer excitatory neurons, significant expansion of oligodendrocytes and immature astrocytic populations, and a unique neuronal subpopulation harboring dysmorphic neurons. Furthermore, we uncovered activated microglia subsets, particularly in FCD IIb cases. This comprehensive study unveils neuronal and glial cell states driving FCD development and epileptogenicity, enhancing our understanding of FCD and offering new directions for targeted therapy development.

## INTRODUCTION

Focal Cortical Dysplasia (FCD) is a malformation of the cerebral cortex and a major cause of drug-resistant epilepsy in children. Treatment of this neurological condition with current anti-seizure medications is often ineffective, frequently requiring surgical removal of the affected brain tissue. FCD is stratified based on neuroimaging, histopathology, and genetic characteristics^1,2^. FCD type II, the most common presentation, is characterized by cytoarchitectural cortical abnormalities including loss of cortical lamination, blurring of the gray-white matter junction, and abnormal cellular development^2^. FCD II cortical lesions can be further divided based on cytological features: FCD IIa is characterized by the presence of dysmorphic neurons, while FCD IIb presents with both dysmorphic neurons and balloon cells. Dysmorphic neurons are abnormal cells featuring enlarged soma and nuclei^1,2^, along with expression of neurofilament markers typical of immature neurons in the developing cortex. On the other hand, balloon cells are characterized by large cell bodies with vitreous, opalescent, and eosinophilic cytoplasm^1,2^. These cells express markers commonly associated with neuroglial progenitors, including nestin and vimentin, indicating their immature nature^3^. High expression of other progenitor cell markers such as c-Myc, SOX2, and Oct-4 were also observed throughout FCD II lesions^4,5^.

At the molecular level, FCD II cases display activation of the mammalian target of rapamycin complex 1 (mTORC1) signaling pathway, which is implicated in cell growth, increased synaptic transmission and plasticity, as well as regulation of immune system responses^6–8^. Hyperactivation of mTOR signaling, which operates through the mTORC1 and mTORC2 complexes, is particularly observed in abnormal cells such as dysmorphic neurons and balloon cells^9^. Increased phosphorylation of the mTORC1 substrates S6K1 and S6 proteins can be used as a biomarker to distinguish FCD type II lesions^4^. Genetic alterations such as somatic gain-of-function mutations in mTOR activating genes and germline loss-of-function mutations of mTOR inhibitors are also found in FCD II^9–11^.

The cellular and epileptogenic mechanisms of FCD are not yet fully understood. Dysmorphic neurons appear to play a pivotal role in generating epileptic discharges^11^, while balloon cells are considered electrically quiescent and may even have an anti-seizure effect^3^. A recent study showed that dysmorphic neurons derive from excitatory neurons while balloon cells have astrocytic lineage^12^. We and other groups have shown that FCD IIb lesions are also characterized by increased microglia and astrocyte reactivity when compared to FCD IIa or healthy controls^13–15^. However, little is known regarding the cellular landscape of FCD lesions and the role of the diverse cortical cell types in the disease.

Single-cell RNA and transposase-accessible chromatin (ATAC) sequencing have been applied to resolve the cellular diversity of the human brain in healthy and disease contexts^16–18^ and to discover cell states associated with neurological conditions^19–21^. Recent multimodal single-cell approaches can obtain multiple measurements from the same cells simultaneously, enabling the integration of epigenetic and transcriptomic modalities to identify cell types involved in diseases^22,23^.

In this study, we applied multi-omics single-cell profiling to characterize the chromatin accessibility and gene expression landscapes in FCD type II, generating a comprehensive multimodal single-cell dataset comprising 61,525 cells from 11 clinical samples of FCD lesions and controls obtained from epilepsy surgeries. Integrative analyses of this dataset unraveled cellular unbalances and cell states underlying FCD pathogenesis, and highlighted neuronal populations susceptible in the disease.

## RESULTS

### High-resolution characterization of the chromatin accessibility and transcriptome landscapes of FCD type II

We performed multi-omics single-cell sequencing of cortical brain tissue from FCD IIa (n=3) and IIb (n=6) lesions, as well as from non-lesion tissues selected as controls (n=2; Figure 1A). Specimens were collected during epilepsy surgery to treat drug-resistant FCD. Internal controls correspond to histologically normal cortical tissues where neither abnormal cells (dysmorphic neurons and balloon cells) nor architectural changes were observed. Dissociated nuclei were profiled using the 10X Genomics Multiome ATAC + Gene Expression assay for simultaneous measurement of chromatin accessibility and gene expression in individual nuclei. Quality control was performed to remove low-quality nuclei and to filter out potential doublets, and Harmony^24^ was applied to ATAC and RNA datasets to integrate samples from different sequencing batches (Figure 1B). The Weighted Nearest Neighbor (WNN) approach^25^ from Seurat was used to create an integrated multimodal (ATAC + RNA) visualization (Figure 1B). The resulting multimodal single-cell compedium comprised 61,525 nuclei with paired chromatin and gene expression measurements. Clustering analysis identified a total of 40 populations (Figure S1A), which were annotated using Azimuth^25^ followed by manual validation using canonical marker genes to define consensus cell types. Azimuth was able to assign labels with high confidence to the majority of clusters (Figure S1B), and none of the clusters were exclusive to a unique sample (Figure S1C). Consensus cell type annotation was performed using a broad cell types (Figure 1C) as well as a higher-resolution taxonomy including neuronal subtypes and cortical layers (Figure 1D).

**Figure 1.**
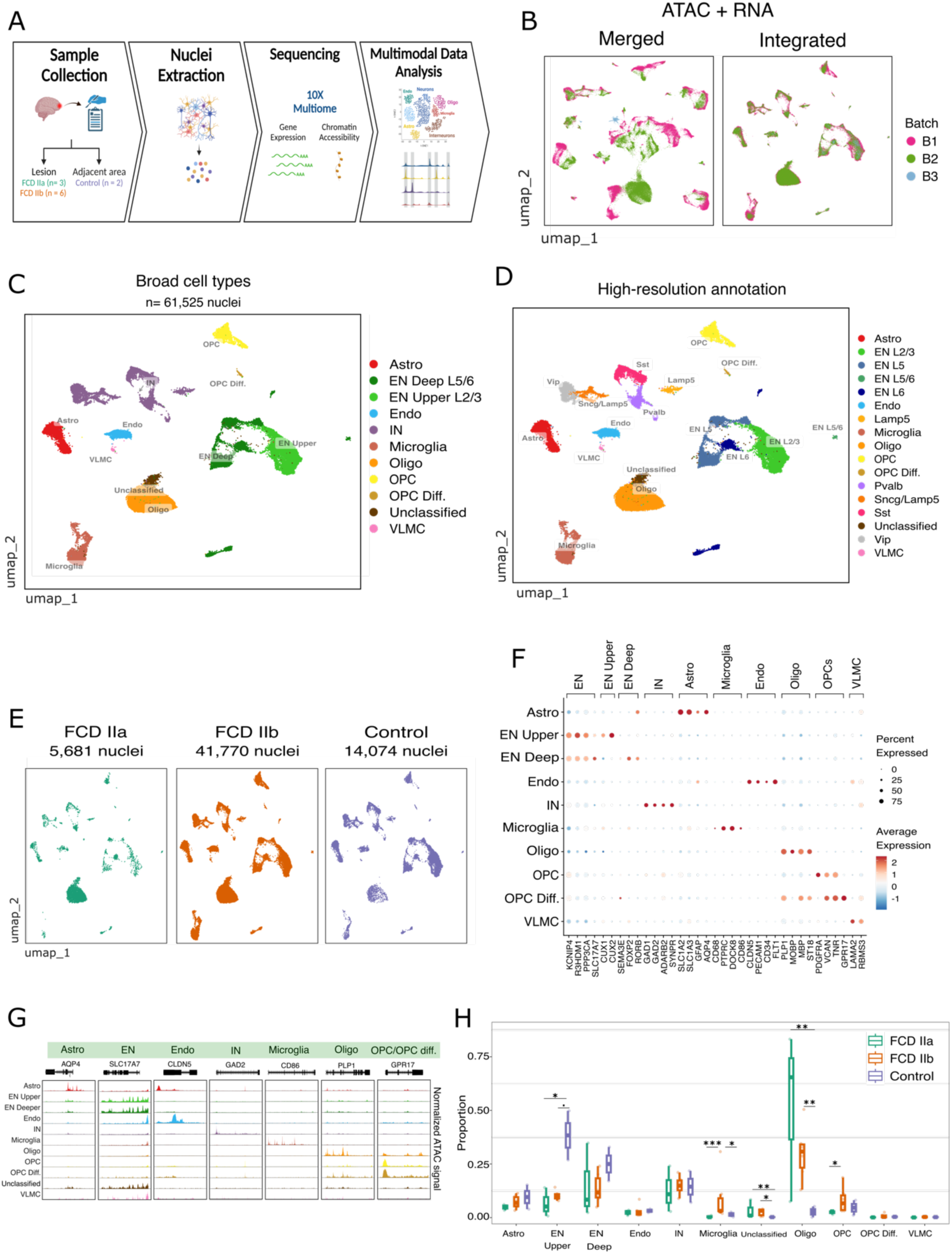
Overview of the multimodal single-cell sequencing in FCD type II. **(A)** Schematic of FCD samples and study design. Cortical tissues from lesions and adjacent non-lesion areas in individuals with FCD IIa and IIb were profiled by multiome single-cell ATAC and RNA sequencing. Created with Biorender.com. **(B)** Uniform manifold approximation and projection (UMAP) visualization of joint modalities (ATAC + RNA) after sample integration. Nuclei are colored by sequencing batch. **(C)** UMAP joint visualization of nuclei colored by broad cell type annotation. **(D)** UMAP joint visualization colored using a high-resolution annotation with classification of neuronal subtypes. **(E)** UMAP plots depicting nuclei by tissue condition. **(F)** Dot plot showing gene expression of known marker genes for major cortical cell types. Dot size corresponds to the fraction of cells expressing the marker, and color corresponds to the expression levels. **(G)** Coverage plots of marker genes depicting chromatin accessibility across cell types. The normalized ATAC signal is depicted in a region of +/- 1kb around gene coordinates. **(H)** Box plots showing cell type proportion by tissue condition. In box plots, the median is indicated by the center line; box limits represent upper and lower quartiles; and whiskers extend to 1.5 times the interquartile range. Overlay dots represent cell type proportions in individual samples. Significant changes in cell types were detected using Speckle. P < 0.1, *P < 0.5, **P < 0.01, ***P < 0.001.

Overall, our annotation identified the various cell types expected in the human cortex such as excitatory and inhibitory neurons, astrocytes, oligodendrocytes, oligodendrocytes progenitor cells (OPC), microglia, endothelial cells, and vascular and leptomeningeal cells (VLMC, Figure 1C). We also identified the various subtypes of excitatory neurons (EN, Figure 1D) including upper layers EN L2/3 (termed EN Upper), and deep layer subtypes EN L5, EN L5/6, EN L6 (EN Deep), while inhibitory neurons (IN) were further classified into Vip, Sst, Pvalb, Lamp5 and Sncg/Lamp5 subtypes (Figure 1D). Based on the histological assessment, most of the sequenced nuclei derived from IIb lesions, followed by internal controls and IIa lesions (Figure 1E). Gene expression and open chromatin accessibility of marker genes were inspected to confirm cell type identity (Figures 1F-G). For instance, *AQP4* (astrocytes), *SLC17A7* (EN), *GAD2* (IN), *PLP1* (oligodendrocytes), *CD86* (microglia), and *GPR17* (OPC) were marker genes with concordant expression and chromatin accessibility in their corresponding cell types.

Using signature enrichment analysis, mTORC1 signaling was found to be elevated in dysplastic tissue across all cell types except VLMC and endothelia (Figure S2A), and cells with the highest mTORC1 activation were identified in FCD IIb (Figure S2B). Thus, multimodal single-cell profiling uncovered the heterogenous composition of FCD type II lesions and recapitulated key aspects of their pathological state.

### Unraveling cellular changes in FCD lesions

Next, we determined cell type abundance changes between FCD and control samples with Speckle^26^, using an approach that accounts for variability in cellular frequencies among patients. The analysis uncovered alterations affecting several broad cell types, including decreased frequency of the EN upper population and increased frequency of oligodendrocytes in both FCD IIa and IIb lesions (Figure 1H). In addition, we found an expansion of the microglia population specifically in IIb donors.

We also leveraged the hierarchical nature of the Azimuth annotation, which classifies excitatory and inhibitory neurons into 116 subclasses based on their expression of markers, to find out how these neuronal subclasses are affected in the disease. The Exc L2 *LINC00507*/*GLRA3*, the most common EN subclass identified in the cortex of FCD samples, was also the most affected EN displaying both the highest number of transcriptional changes and frequency reduction in lesions (Figure 2A,B). Among IN, FCD IIa displayed an increased frequency of Parvalbumin-expressing (Pvalb) interneurons Inh L3 Pvalb/*SAMD13* (Figure S3C); yet, this was not the subclass with most transcriptional changes (Figure 2D). On the other hand, somatostatin-expressing (Sst) subclass Inh L1-2 Sst/PRRT4 and PAX6-expressing subclass Inh L1 PAX6/*MIR101-1* were affected only in FCD IIb when compared to controls (Figure 2C). Altogether, this data shows that cellular abundance changes affect both neuronal and non-neuronal populations in FCD pathogenesis, and revealed the vulnerability of excitatory neurons in the disease.

**Figure 2.**
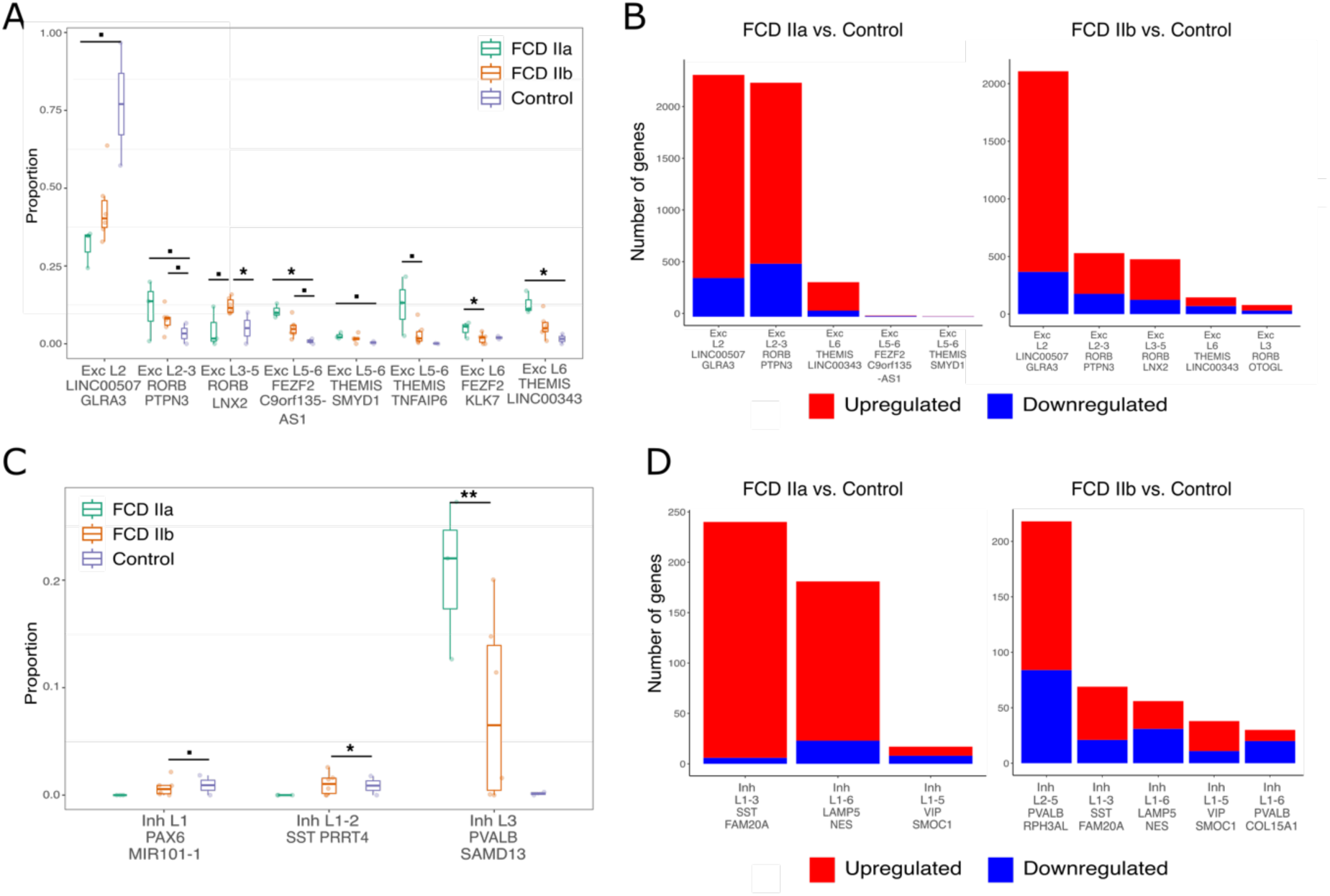
Excitatory and inhibitory neuron subtypes affected in FCD. **(A)** Box plots showing excitatory neurons (EN) subtype frequency in FCD and controls. In box plots, the median is indicated by the center line; box limits represent upper and lower quartiles; and whiskers extend to 1.5 times the interquartile range. Overlay dots represent proportions in individual samples. Significant changes were detected using Speckle. P < 0.1, *P < 0.5, **P < 0.01, ***P < 0.001. **(B)** Bar graphs depicting the number of differentially expressed genes (DEGs) in EN subtypes. DEGs between FCD and controls were computed using MAST with adj. P < 0.05. Only the five subtypes with the most transcriptional changes are shown. **(C)** Box plots showing inhibitory neurons (IN) subtype frequency in FCD and controls. Box plots legends and significance levels are defined as in (A). **(D)** Bar graphs depicting DEGs in IN subtypes. DEGs between FCD and controls were computed using MAST with adj. P < 0.05. Only the five subtypes with the most transcriptional changes are shown.

### Identification of a disease-specific population containing dysmorphic neurons

During annotation, we found a heterogeneous cluster containing both excitatory neurons and oligodendrocytes that we initially termed Unclassified since Azimuth could not confidently assign a cell type (Figure 1C). This cluster was highly disease-specific as it contained 1,308 cells, being that 1,285 cells (98%) originated from FCD (Figure 3A). Despite clustering next to oligodendrocytes after data integration, 54% of nuclei were classified as EN and 29% as oligodendrocytes. The cluster was detected in all 9 FCD samples, denoting its reproducibility across donors (cluster #15, Figure S1C). To further characterize this disease-specific (DS) population, we identified its markers and performed pathway enrichment analysis. Notably, all top enriched pathways were related to neurodegenerative diseases including Parkinson’s, amyotrophic lateral sclerosis, and Alzheimer’s (Figure 3B). Also, the majority of marker genes were associated with neuronal processes including oxidative phosphorylation, Ca^2+^ transport, neuron development, and neurofilament subunits (Figure 3C).

**Figure 3.**
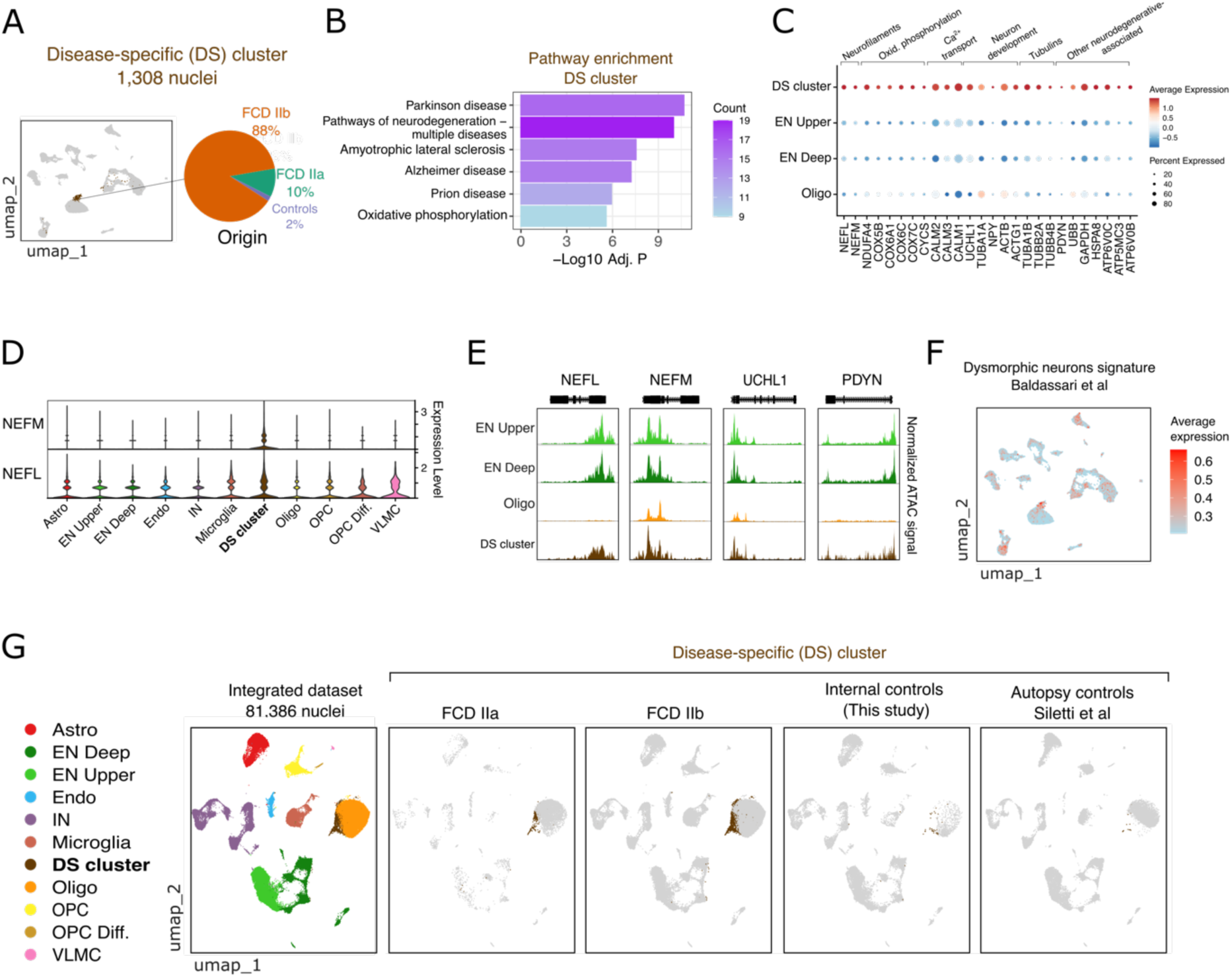
Characterization of a disease-specific cluster containing dysmorphic neurons. **(A)** UMAP highlighting the disease-specific (DS) cluster. The pie chart indicates the percentage of nuclei from FCD IIa, IIb, and controls within the cluster. **(B)** Pathway enrichment of DS cluster markers. Marker genes were identified using Seurat’s FindMarkers and KEGG pathway enrichment was performed using g:Profiler. The bars represent the significance score of pathway enrichment, and the color indicates the number of genes in each pathway. **(C)** Dot plot denoting expression of marker genes associated with neurons in the DS cluster. Neuronal genes are grouped by biological function. Dot size and color correspond to expression frequency and level in the cluster, respectively. **(D)** Violin plots denoting gene expression of *NEFM* and *NEFL* neurofilaments in the various cell types. **(E)** Coverage plots of neuronal genes expressed in the DS cluster. The normalized ATAC signal is depicted in a region of +/- 1kb around gene coordinates. **(F)** UMAP showing the average expression of a dysmorphic neuron (DN) signature obtained from Baldassari et al. Signature expression was calculated using the AddModuleScore function from Seurat. **(G)** UMAP RNA plots showing the DS cluster population in FCD IIa and IIb lesions, histologically normal tissue from FCD individuals (internal controls), and healthy cortex (autopsy controls). The leftmost UMAP depicts annotated cell types in the integrated dataset, which was created by integrating scRNA-seq data from this study with Siletti et al using Harmony.

In particular, we found two neurofilament subunits *NEFL* and *NEFM* overexpressed in the DS cluster (Figure 3C). *NEFM* was exclusively expressed in the DS cluster and not detected in any other cell type in either lesions or non-lesion tissues (Figure 3D). The chromatin accessibility in the *NEFM* and *NEFL* loci corroborates with their active gene expression in the DS cluster (Figure 3E).

Antibodies against neurofilaments are often used to identify dysmorphic neurons in dysplastic tissue during histopathological diagnosis of FCD^2,11^. Based on this observation, we used a signature enrichment analysis to inspect whether the DS cluster harbor dysmorphic neurons. Specifically, we analyzed a 30-gene signature expressed in a purified dysmorphic neuron population selected using laser capture microdissection in FCD tissue^27^. We found that nuclei with the highest expression of this signature were overlapping the DS cluster (Figure 1F). Finally, to find out whether the DS cluster could be detected in healthy individuals, we integrated our FCD dataset with single-cell RNA-seq from cortical samples obtained from age-matched autopsy controls^17^. This analysis confirmed that the DS cluster was not present in the normal healthy cortex (Figure 1G). Taken together, these data demonstrate that the DS population is uniquely present in FCD lesions and comprises dysmorphic neurons expressing genes associated with neurodegenerative conditions.

### Pathological microglia cell states in FCD IIb

To further characterize microglia states in FCD, we performed subclustering and re-annotation of the broad microglial compartment. Subclustering and marker detection with Seurat uncovered 7 subpopulations with clearly distinct gene signatures (Figure S3; full list of subcluster markers provided in Supplemental Table 2). Diverse microglia signatures identified by single-cell sequencing have been previously reported in neurological diseases^20,21,28^. We used signatures of microglia subsets derived from healthy and epileptic temporal cortex^21^ to annotate subclusters in FCD. Using this approach, we identified three distinct microglia states in FCD lesions (Figure 4A): a homeostatic subset (the largest population corresponding to subcluster 0); a subset expressing high levels of *CD74* (subcluster 1, *CD74*^+,^), and a subset expressing high levels of *CD83* (subclusters 2 and 5, *CD83*^+^). We also identified small subclusters of lymphocyte/T cells, B cells, and perivascular macrophages that were annotated using classical lymphocyte markers (Figure 4A). As shown in Figure 4B, *CD74*^+^ microglia expressed marker genes related to antigen processing and presentation/MHC class II genes (*HLA-DRA*, *HLA-DPA1*, *HLA-DPB1*, *HLA-DRB1*, *HLA-DQA1*), adaptive immune system response and microglial cell activation (*CD74*, *SC1N*, *C1QA*, *C1QB*, *C1QC*), and interferon signaling (*IFNGR1*, *JAK2*, *B2M*). On the other hand, *CD83*^+^ microglia expressed high levels of pro-inflammatory cytokines including *IL1B*, *CCL2*, *CCL3*, *CCL4*, and *TNFSF18*. Interestingly, *CD83* and *CD74* expression are largely non-overlapping, indicating that these surface markers can be effectively used to distinguish microglia states in FCD (Figure 4C).

**Figure 4.**
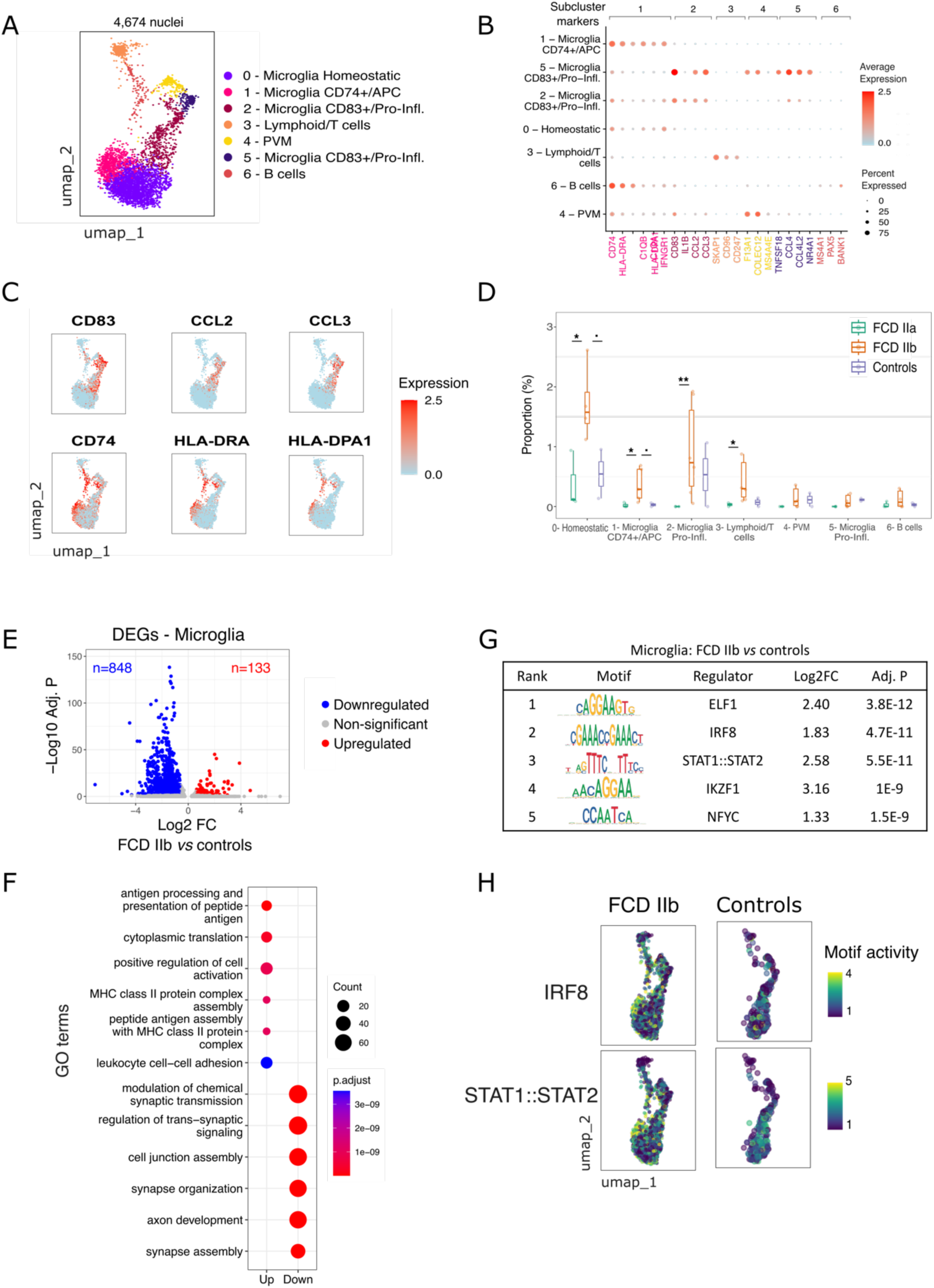
Microglia pathological cell states and activation in FCD type IIb. **(A)** UMAP visualization of microglia and lymphoid subclusters sequenced in FCD lesions. **(B)** Feature plots depicting expression of key genes distinguishing microglia subsets *CD83*^+^ (*CD83*, *CCL2*, *CCL3*) and *CD74*^+^ (*CD74*, *HLA-DRA*, *HLA-DPA1*). **(C)** Dot plot denoting the expression of key marker genes in microglia and lymphoid subclusters. **(D)** Box plots denoting subcluster frequency by tissue condition. Significant changes were detected using Speckle. Box plots and significance legends are defined in Fig. 1H. **(E)** Volcano plot depicting differentially expressed genes (DEGs) in FCD IIb microglia. DEGs were obtained using a Wilcox test after SCT transformation with correction for sequencing batch, mitochondrial gene percentage, and brain region. The *x*-axis is the fold-change (log2) expression in FCD IIb *vs* controls, and the *y*-axis depicts the significance of the change (-log10 of the adjusted P-value). The color indicates the gene status. **(F)** Dot plots showing top enriched GO terms in up- and down-regulated genes from (E). Dot size and color correspond to the number of genes associated with the GO term and enrichment P-value, respectively. **(G)** Top five differential motifs in microglia from FCD IIb versus controls, as measured by chromVAR (see Methods). **(H)** Feature plots depicting IRF8 and STAT1/STAT2 motif activity in microglia open chromatin regions from FCD IIb and controls. Cells are colored by the chromVAR score.

Of note, homeostatic microglia and activated states are expanded only in FCD IIb tissues (Figure 4D). Compared to controls, FCD IIb had a higher frequency of *CD74*^+^ and homeostatic subclusters. Compared to FCD IIa, the expansion included *CD74*^+^ and *CD83*^+^ subpopulations as well as lymphoid/T cells (Figure 4D).

Next, we performed differential expression analysis and identified 133 upregulated genes in FCD IIb microglia compared to controls (Figure 4E). GO terms associated with upregulated genes included antigen processing and presentation, MHC class II protein complex, and positive regulation of cell activation (Figure 4F), in line with the expansion of *CD74*^+^ microglia in FCD IIb. Motif activity analysis using scATAC-seq uncovered that top upregulated motifs in FCD IIb microglia included IRF8 and STAT1/STAT2 (Figure 4G), which are regulators related to microglia activation. Cells with the highest IRF8 and STAT1/STAT2 motif activity overlapped the *CD74*^+^ microglia (Figure 4H). Taken together, these data highlight the emergence of pathological microglial states is specific to FCD IIb, and indicated that microglia activation in FCD IIb is mediated by the *CD74*^+^ subpopulation.

### Chromatin changes and cell-type specific gene regulation in FCD

Next, we aimed to find open chromatin regions associated with gene regulation in FCD. First, we found the set of differentially accessible chromatin regions (DACRs), which are peaks with gain or loss of chromatin accessibility, by comparing chromatin profiles across cell types in FCD against controls. Overall, FCD IIa had a higher number of DACRs, especially in upper and deep excitatory neurons (Figure 5A). Most of the DACRs in FCD IIa displayed a gain of accessibility in the lesion (opening peaks), while DACRs in FCD IIb were characterized by a loss of chromatin accessibility (Figure 5A). Further analysis using Encode annotation found that most DACRs were located within enhancer regions, with 34% overlapping proximal enhancers near a transcriptional start site and 44% overlapping distal enhancers (Figure 5B).

**Figure 5.**
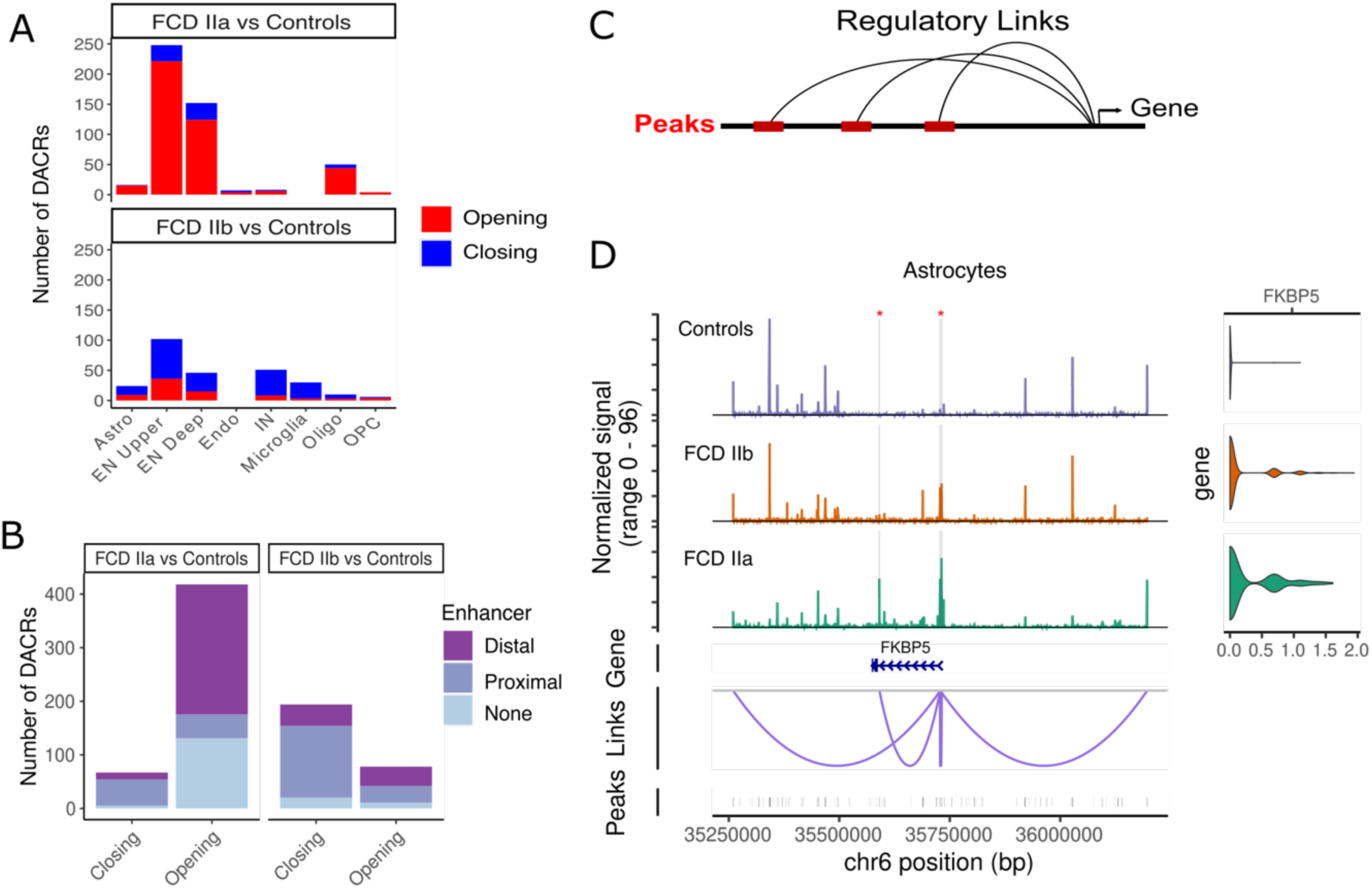
Open chromatin changes and cell-type specific regulatory links in FCD. **(A)** Barplots depicting differentially accessible chromatin regions (DACRs) between FCD tissues and controls. The number of chromatin regions with opening or closing status (gain or loss of chromatin accessibility, respectively) is indicated for each cell type. **(B)** Enhancer types overlapping DACRs. The number of opening and closing peaks associated with distal or proximal enhancers from Encode are indicated. **(C)** Diagram depicting the inference of regulatory links identifying neighboring open chromatin regions (ATAC peaks) associated with gene expression using linear regression (see Methods). **(D)** Regulatory links predicted for *FKBP5* in astrocytes. ATAC coverage tracks and corresponding expression levels in controls, FCD IIb, and IIa tissues are depicted at the top. Regions highlighted in gray indicate opening peaks in lesions and red asterisks indicate Encode-annotated enhancers. The gene track shows the location of the *FKBP5* locus in chromosome 6, and the bottom track shows all peaks contained in the chromosomal region.

We then sought to understand whether these DACRs were driving gene regulation. To achieve this, we implemented a machine-learning approach based on random forests to learn regulatory links using the paired single-cell chromatin and gene expression readouts in the dataset (see Methods, Figure 5C). The method revealed open chromatin regions most likely to predict gene expression in each cell type. For instance, we identified regulatory links for *FKBP* prolyl isomerase 5 (*FKBP5*), a gene within the mTORC1 pathway upregulated in FCD astrocytes (Figure 5D). The regulatory links involved DACRs with gain of chromatin accessibility in FCD (regions highlighted in Figure 5D), suggesting that the increased *FKBP5* expression is accompanied by chromatin opening. These peaks are also putative *FKBP5* cis-regulatory elements since they are located in regions with enhancer activity according to Encode. Similarly, we also found opening chromatin peaks located in enhancers linked to *C1QA* and *C1QB*, which were genes involved in microglia cell activation increased in FCD IIb (Figure S4). Taken together, these analyses demonstrated that chromatin changes affecting enhancer regions are likely driving cell type-specific gene regulation in FCD.

### Immature astrocyte cell states and balloon cells in FCD

We explored whether the astrocytic population is affected in FCD by employing an unsupervised trajectory analysis. The trajectory organized astrocytes according to their differentiation status, with immature and mature astrocytes on opposite ends of the trajectory (Figure 6A). The root of the trajectory was chosen based on the expression of immature astrocyte markers (Figure 6B). Remarkably, the quantification of FCD and control cells along the trajectory revealed an expansion of immature astrocytes accompanied by a decrease of mature astrocytes in FCD tissues (Figure 6C). On the other hand, fully mature astrocytes were highly represented in controls (Figure 6C). We also analyzed the expression of gene signatures corresponding to balloon cells, which are known to originate from the astrocytic lineage^27^ as well as reactive astrocytes often associated to neurodegenerative diseases^19^. Scores for both balloon cells and reactive astrocytes signatures were more elevated in astrocytes from FCD lesions and overlapped with immature states at the beginning of the trajectory (Figure 6D, top). Of note, immature astrocytes identified as balloon cells were mostly exclusive to FCD IIb cases, while reactive astrocytes were present in both FCD subtypes (Figure 6D, bottom). Genes associated with balloon cells and reactive astrocytes including *CHI3L1*, *HSPB1*, *SERPINA3*, *GFAP*, and *VIM* were up-regulated in FCD IIb astrocytes compared to FCD IIa or internal controls (Figure 6E). Taken together, these analyses showed that immature astrocyte cell states, including balloon cells and reactive astrocytes, are expanded in the disease, indicating that normal astrocyte differentiation is impaired in FCD.

**Figure 6.**
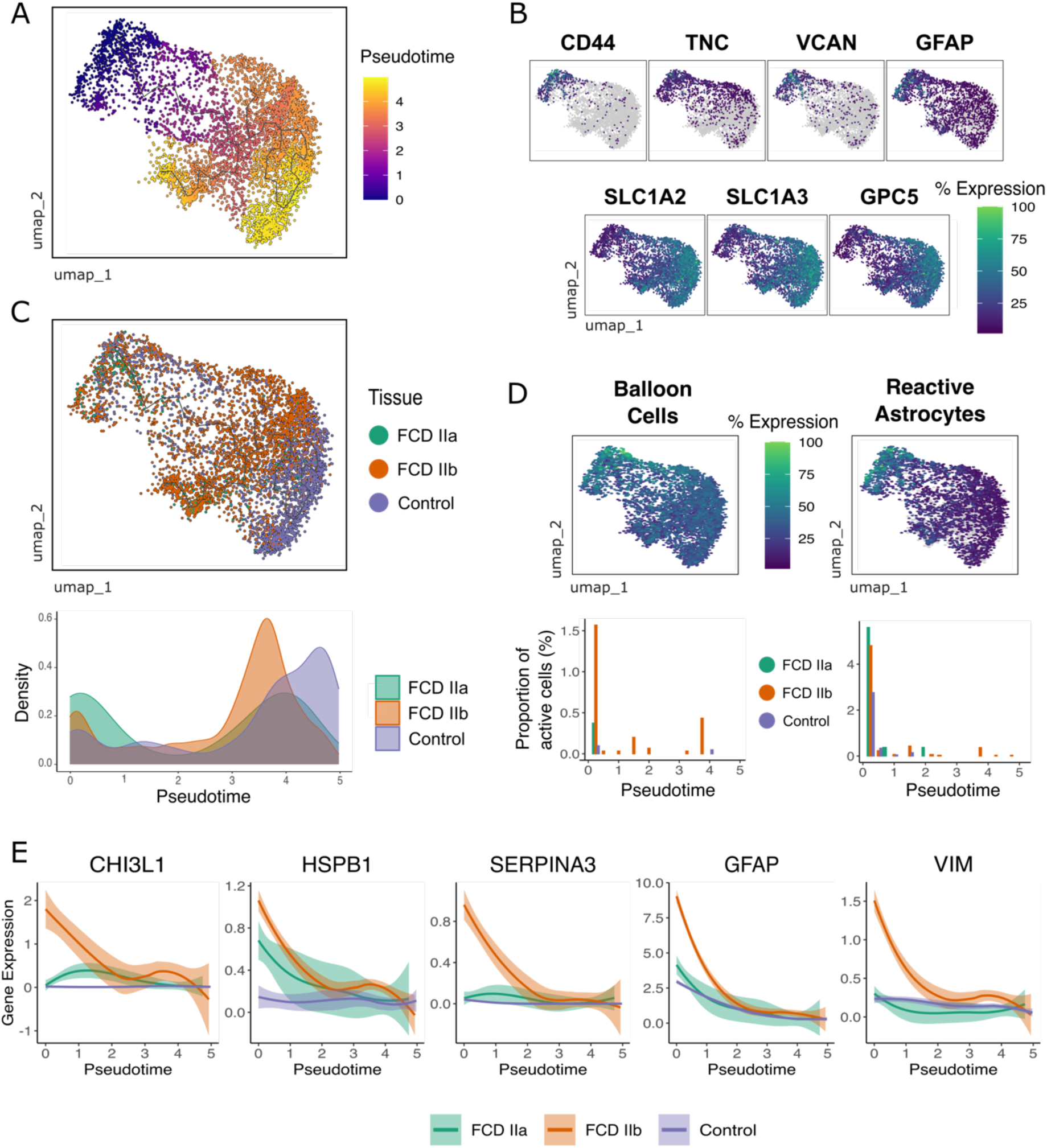
Impaired Astrocyte Differentiation in FCD. **(A)** UMAP visualization of astrocytes temporally ordered on a trajectory as a function of pseudotime built by Monocle3. **(B)** Feature plots depicting gene expression of immature astrocyte markers *CD44*, *TNC*, *VCAN*, and *GFAP* (top), and differentiated astrocyte markers *SLC1A2*, *SLC1A3*, and *GPC5* (bottom). **(C)** UMAP visualization and density quantification along the trajectory. Density plots indicate the normalized density of cells along the pseudotime according to tissue condition. **(D)** Feature plots depicting expression of balloon cells and reactive astrocytes gene signatures computed by AddModuleScore from Seurat (top). Bar graphs (bottom) quantify the proportion of active cells for these signatures along the pseudotime trajectory. Active cells were defined using the AUCell method and are colored by tissue condition. **(E)** Expression levels of genes associated with balloon cells and reactive astrocytes *CHI3L1*, *HSPB1*, *SERPINA3*, *GFAP*, and *VIM* in the trajectory and by tissue condition.

## DISCUSSION

FCD is a leading cause of drug-resistant epilepsy in children and adolescents, accounting for up to 50% of epilepsy surgeries in this age group^10^. This neurological condition was found to be the most common diagnosis among children with drug-resistant focal epilepsy requiring surgical intervention^29^. Despite advances in the genetic characterization of FCD, brain surgery remains the only treatment option to reduce seizures in refractory cases. Our limited understanding of the molecular mechanisms and the role of diverse brain cell types in FCD epileptogenicity is a major obstacle to developing targeted treatments. To address this need, we employed a cutting-edge multi-omics single-cell profiling approach to create a high-resolution cellular census of the human cortex affected by FCD. We leveraged this unique dataset to uncover cellular changes and coordinated chromatin and transcriptional alterations in both neuronal and glial populations in response to the disease. Overall, we uncovered neuronal, microglial and astrocytic pathological cell states involved the disease.

First, our data showed that both excitatory and inhibitory neurons undergo changes in cellular abundance and transcriptional profiles during FCD development, yet neuronal loss affects mostly the EN L2/3 subtype. Interestingly, EN in upper cortical layers were also found to be reduced in cortical samples of patients with multiple sclerosis^30^ and temporal lobe epilepsy^31^, highlighting their vulnerability not only in FCD but in other neurological conditions.

In addition, we characterized a neuronal population specific to FCD lesions expressing a dysmorphic neuron-signature and neurofilament genes. This neuronal population was notably absent in non-lesion controls and the healthy cortex from autopsy cases. Also, the expression of *NEFM*, encoding the medium polypeptide subunit of the neurofilament protein, was found to be restricted to this subpopulation. Neurofilaments detected in the blood have emerged as reliable diagnostic biomarkers in several neurological diseases^33^. The remarkable specificity of *NEFM* expression in dysplastic cells suggests its potential as a blood biomarker for FCD, prompting the need for additional research to explore its utility as a molecular diagnostic tool in the disease.

Furthermore, we described the expansion of microglia and the emergence of pathological cell states specifically in FCD IIb lesions. We identified two FCD-associated microglial subpopulations with distinct signatures: a *CD74*^+^ subset overexpressing antigen presentation molecules (MHC class-II components), and a *CD83*^+^ subset with increased expression of pro-inflammatory cytokines. Previous scRNA-seq studies have identified microglial populations in neurological conditions^20,21,28^. For instance, using scRNA-seq followed by in situ immunohistochemistry, Olah et al^21^ described the existence of nine microglial subpopulations in the human cortex (prefrontal cortex of autopsy samples and temporal cortex of epilepsy), including the *CD83*^high^ and *CD74*^high^ subsets. Mathys et al^34^ also identified a microglia subset with increased expression of *CD74* and MHCII-related genes in patients with Alzheimer’s disease. A *CD74* subset was also described in the pathology of multiple sclerosis^35^, and *CD74* has been proposed as a marker for reactive microglia due to its increased expression levels in disease^36^. Although there is no consensus in the field on how to annotate microglia cell states, we adopted the terminology from Olah et al since the *CD83* and *CD74* molecules were identified as markers of microglia subclusters in FCD. Of note, we also found higher lymphoid/T-cells infiltration is specific to FCD IIb, in agreement with previous experimental findings^14^.

There is strong emerging evidence of dysregulated neuroinflammation mediated by microglia in FCD and epilepsy. Zimmer et al^14^ showed that FCD IIb elicits a stronger inflammatory response characterized by increased expression of the *HLA-II* and *CCL2* in microglia near balloon cells. Kumar et al^37^ described immune cell infiltration and increased expression of pro-inflammatory genes (e.g. *IL1B*, *CCL4*, *CCL2*) in microglia from epilepsy samples. Therefore, we postulate that the CD74^+^ subset (expressing antigen presentation molecules) and the CD83^+^ subset (expressing pro-inflammatory cytokines) identified in this study are the microglial populations mediating immune system activation and neuroinflammation in FCD IIb.

Finally, we showed that astrocyte differentiation is impaired in FCD, and immature astrocytic cell states are expanded in lesions. This shift from mature to immature astrocytes can be explained in part by the appearance of reactive astrocytes and balloon cells, which are abnormal cell populations typically found in the lesion microenvironment, that we characterized as immature astrocytic states. Specifically, pseudotime trajectory analysis revealed that reactive astrocytes resemble immature-like cells that emerge in FCD lesions, supporting the view that astrogliosis induces a partial reversion of astrocyte maturation^38^. Similarly, we showed that balloon cells also represent immature astrocytic cell states. Balloon cells were identified exclusively in FCD IIb, consistent with the histological characterization of the clinical samples. Moving forward, it will be important to understand the contribution of these immature astrocytes to FCD. Mature astrocytes are known to maintain neuronal homeostasis and prevent excessive neuronal activity through neurotransmitter reuptake^38^, and their dysregulation can lead to epileptogenesis^39^. Thus, these immature astrocytic populations might add to the intrinsic epileptogenicity in FCD.

Paired multimodal single-cell experiments enable exploration of cell-type specific gene regulation by learning the relationship between chromatin accessibility and transcription in the same cell^40^. We explored the integration of these paired readouts by predicting cell-type specific regulatory links and identifying enhancer regions controlling key genes in the mTORC1 pathway and microglia activation. Indeed, we anticipate that the high-resolution multimodal dataset produced herein will be useful in future studies integrating chromatin and transcriptomic data, including unraveling the impact of non-coding variants in open chromatin regions in gene expression or identifying chromatin enhancers driving gene dysregulation in FCD.

The findings presented in this study offer promising new directions for the development of targeted drug treatments for FCD. By elucidating the underlying molecular and cellular mechanisms common and specific to FCD IIa and IIb at single-cell resolution, this research provides valuable insights that could inform the development of tailored therapeutic strategies for each disease subtype. For instance, mTORC1 signaling was found to be elevated in FCD in most cell types. Thus, mTOR inhibitors or antisense oligonucleotides targeting the mTOR pathway hyperactivation may represent promising strategies for targeted treatment of FCD^11^. Gene therapies targeting the disease-specific population containing dysmorphic neurons can also be explored since dysmorphic neurons are believed to play a crucial role in seizure generation in FCD^3^. Further investigation into marker genes associated with this population may lead to the identification of pharmacological interventions aimed at improving seizure control in the disease.

Additionally, several rare neurological conditions lead to epilepsy^41^. These orphan diseases pose significant challenges to research due to their low prevalence and unresolved pathology^41^. However, the integration of single-cell datasets may provide a powerful approach to uncover pathological cell states shared across various forms of rare focal epilepsies. By comparing cellular profiles, it may be possible to identify common abnormal cell states, molecular signatures and pathways, offering insights into disease mechanisms and potential therapeutic targets applicable to multiple epilepsy subtypes^32^.

In conclusion, our study uncovered diverse cellular abundance changes in FCD and identified key neuronal, microglial and astrocytic subpopulations involved in the disease. These findings improve our understanding of the condition and suggest novel avenues for developing targeted treatments.

## STAR METHODS

### Clinical samples and neuropathological diagnosis

Fresh brain samples were collected from patients who underwent surgery for intractable seizures after clinical diagnosis of FCD at Hospital de Clínicas, University of Campinas. Surgical specimens were (i) formalin-fixed paraffin-embedded (FFPE) or (ii) snap-frozen in liquid nitrogen and stored at −80°C. FFPE samples were submitted to diagnostic routine in serial 4 µm-sections stained with hematoxylin and eosin (H&E) and submitted to immunohistochemical reactions. For the latter protocol, the sections were exposed to antibodies against NeuN (neuronal marker; 1:1,000, clone A60, Merck Millipore, cat#MAB377, Temecula, CA, USA), MAP2 (neuronal marker; 1:1,000, clone M13, Thermo Fisher, cat#13-1500, Waltham, MA, USA), SMI 32 (neuronal marker; 1:2,500, clone SMI 32, Biolegend, cat#SMI-32R, San Diego, CA, USA), GFAP (astrocytic marker, 1:100, clone 6F2, Dako/Agilent, cat#M0761, Santa Clara, CA, USA), vimentin (1:100, clone V9, e-Bioscience/Thermo Fisher, cat#14-9897-82, Waltham, MA, USA), CD34 (1:50, clone QBEnd-10, Dako, cat#M7165, Glostrup, Denmark) and CNPase (myeloarchitecture marker; 1:500, clone 11-5 B, Millipore, cat#MAB326, Darmstadt, Germany), for 18 h at 4°C. Then, a detection solution containing the secondary antibody and peroxidase (AdvanceTMHRP®, Dako, cat#K4068, Glostrup, Denmark; or EnvisionTM Flex+, Dako, cat#K8002, Glostrup, Denmark) was added for 30 min at 37°C. 3,3-diaminobenzidine (DAB) was used as a chromogenic substrate and counterstaining was performed with hematoxylin. Negative controls (without primary antibody) were run concurrently with all immunohistochemical reactions.

Samples were classified as FCD IIa or IIb according to the guidelines of the International League Against Epilepsy (ILAE)^2^. Specifically, specimens exhibiting cortical dyslamination, hypertrophic and dysmorphic neurons [disoriented neurons with anomalous cytoplasmic distribution of Nissl substance and accumulation of non-phosphorylated neurofilament (SMI 32-positive)] without or with balloon cells (large cells with opaque eosinophilic cytoplasm, vesicular nucleus and immunopositivity for vimentin) were classified as FCD ILAE type IIa or IIb, respectively. Internal controls were derived from histologically normal tissue where neither abnormal cells nor architectural changes were observed. Detailed patient information is provided in the Supplemental Table 1. All procedures were approved by the University of Campinas’s Research Ethics Board, and written informed consent was obtained from all patients before surgery.

### Nuclei isolation and multiome library construction

The nuclei extraction protocol for human brain tissue was adapted from Jessa et al^42^. Briefly, frozen tissue was dounce homogenized and incubated for 2 minutes in lysis buffer containing 0.1% NP-40. Then, suspension was filtered through a 30 µm cell strainer (MACS), and washed in buffer containing 10% BSA to remove debris, followed by a centrifugation at 500g and 4°C for 5 minutes. Next, the suspension was carefully layered on double 29% and 50% iodixanol (Sigma) cushion layers, and centrifuged at 10,000g and 4°C for 30 minutes for additional removal of debris and cell aggregates. Retrieved nuclei were resuspended and counted. Nuclei were stained with Trypan blue and counted using a hemocytometer. Nuclei integrity was evaluated under a microscope using 40X or 60X magnifications, and nuclei were considered viable if they were roundish and with an intact membrane. Next, we proceeded to the steps of the library construction, including open chromatin transposition and droplet (GEM) formation using a Chromium Controller equipment (10X Genomics). Libraries were prepared using the Multiome ATAC + Gene Expression kit (10X Genomics) following the manufacturer’s protocol CG000338 revision E. The quality of purified ATAC and RNA libraries was assessed using a TapeStation equipment (Agilent Technologies).

### Sequencing and raw data processing

Multiome libraries were sequenced on either Illumina HiSeqX Ten or Illumina NovaSeq 6000 instruments according to the sequencing depth and the read length recommended in the Multiome kit protocol (CG000338). Demultiplexing, genome alignment, gene quantification, and peak accessibility in single nuclei were performed with the Cell Ranger ARC pipeline^43^. Specifically, fastq files were generated using cellranger mkfastq, then processed using the cellranger-arc count v.2.0.2 and the human genome GRCh38 to generate the count matrices for the RNA and ATAC modalities. Next, we performed data processing, normalization, and batch integration on both assays independently, using Seurat v.5.0.1^44^ and Signac v.1.12.0^45^ in the R environment v.4.1.2.

### scRNA-seq analysis

Samples were preprocessed in Seurat to remove low-quality nuclei using the following criteria: 400 < nFeatures < 7,000, 500 < UMI counts < 50,000, and percentage of mitochondrial reads (percent.mt) < 15. Identification of doublets was performed per sample using scDblFinder v.1.8.0^46^. Next, standard Seurat data processing and normalization steps were performed: NormalizeData, FindVariableFeatures, ScaleData, RunPCA, RunUMAP, FindNeighbors, and FindClusters. For each sample, the filtered counts matrix was log-normalized with regression of sequencing mitochondrial gene percentages. The RunUMAP function was used with the first 40 PCs identified in the elbow plot to perform dimensional reduction. Clustering was performed using FindNeighbors and FindClusters functions using the smart local moving (SLM) algorithm, with a resolution of 1.2. To correct batch effect among samples, we used Harmony v.1.2^24^ to integrate sample-level PCA projections using automatic hyperparameter optimization and default parameters.

Harmony implements an algorithm that projects cells into a common space where they cluster by cell type rather than other sample attributes such as sequencing batch.

Finally, we used SCTransform to create a normalized expression matrix containing all samples, and with regression of sequencing batch, brain region, and mitochondrial gene percentages. DEGs between FCD and controls were calculated using MAST or the Wilcox test implemented in Seurat as indicated. For MAST, DEGs were selected using an adjusted P value of less than 0.05. The Wilcox test was applied using the SCT normalized data, using a log2 fold-change threshold of 0.25 and an adjusted P value of less than 0.05.

### scATAC-seq analysis

Samples were preprocessed in Signac to remove low-quality nuclei using the following criteria: 1,000 < ATAC fragments < α(ATAC fragments), 400 < ATAC fragments in peaks < α (ATAC fragments in peaks), percent.mt < 15, nucleosome signal < 2, TSS enrichment < 2. Upper limits (α) of ATAC fragments per sample were computed as mean plus 2 standard deviations. Peaks within each sample were identified using MACS2^47^. Standard processing and normalization steps were performed as follows: RunTFIDF, FindTopFeatures, RunSVD, and FindClusters. For each sample, the peak counts matrix was normalized using the RunTFIDF function, to correct for differences in nuclei sequencing depth. Dimensional reduction was performed with latent semantic indexing (LSI) using the RunSVD function. UMAP projection was performed utilizing LSI components 2-30, as the first LSI component was confirmed to be a technical variation. Before sample integration, we created a common peak set across samples using MACS2 and quantified peaks in each sample using the FeatureMatrix function. Peaks in non-standard chromosomes and blacklisted regions were removed from downstream analyses. We then re-computed RunTFIDF, FindTopFeatures, and RunSVD on the merged object and applied Harmony to integrate sample-level low-dimensional cell embeddings (LSI components 2-30) using the same parameters defined in the RNA modality. Differentially accessible chromatin regions (DACRs) between FCD tissues and controls were calculated using FindMarkers with the LR test, with a min.pct of 0.05, an adjusted P value less than 0.05, and included the number of ATAC fragments as a latent variable.

### Multimodal data integration

To integrate gene expression data with chromatin accessibility data, we performed Weighted Nearest Neighbor (WNN) analysis^25^, an unsupervised approach that learns the relative utility of each modality and builds a joint graph representing a weighted combination of RNA and ATAC modalities. WNN was performed using the FindMultiModalNeighbors function with 20 neighbors (k.nn), based on the UMAP reductions obtained after Harmony integration. The WNN graph was used for creating a joint UMAP visualization and clustering based on the WNN graph was performed with the FindClusters function using the smart local moving (SLM) algorithm. WNN clusters with fewer than 50 nuclei, a high percentage of doublets, or enriched in mitochondrial markers were removed from downstream analyses.

### Cluster annotation

We used the Azimuth R package^25^ to perform cell type annotation in the FCD dataset, using as a reference the Allen atlas of the human cortex^48^. Azimuth performed label-transferring by assigning to each cell in the FCD dataset the most similar cell type in the reference based on gene expression similarity. The assigned cell type chosen for each cluster was defined as the cell type with the highest mean frequency within the cluster. The Azimuth annotation was manually verified using the marker genes shown in Figure 1C. Cluster and subcluster markers were calculated using the function FindAllMarkers based on the SCT normalized data, with the Wilcox test using a log2 fold-change threshold of 0.25 and an adjusted P value of less than 0.05.

### Differential cellular abundance

We used the function propeller of the Speckle R package^26^ to detect changes in cellular composition between FCD tissues and controls. The propeller function calculated cell type proportions in biological replicates, and fitted a linear model for each cell type with the tissue condition and brain cortical region as covariates.

### Integration with scRNA-seq from normal cortex

We obtained normal cortex scRNA-seq data from three autopsy donors with no history of neurological disease from Silleti et al^17^. To reduce computational costs, we randomly sampled 10,000 neuronal and 10,000 non-neuronal cortical nuclei from these donors, and processed this data along with our FCD dataset following the procedures described in the “scRNA-seq analysis” section. We used Harmony v.1.2 to integrate all samples with automatic hyperparameter optimization and default parameters. Clustering was performed with FindNeighbors and FindClusters functions using the smart local moving (SLM) algorithm and a resolution of 2.0.

### Gene signature analysis

We used the AddModuleScore function from Seurat and the AUCell^49^ R package to calculate gene set enrichment at the nuclei level. The AddModuleScore function from Seurat was calculated based on the SCT-normalized RNA data. AUCell computes the gene set enrichment scores based on ranked gene lists (highest to lowest gene expression) in each cell. AUCell scores were computed using the calcAUC function using the top 5% of genes. The obtained threshold was used to classify cells with active status for the given signature.

### Motif enrichment analysis

We performed motif analysis using chromVAR v. 3.3.2^50^ and Signac. Motif activity at the nuclei level was calculated using the RunChromVAR function in Signac after matching the set of background peaks. Differential motif activity was computed using FindAllMarkers and FindMarkers (pairwise comparisons) using the LR test, with a min.pct of 0.05, an adjusted P value less than 0.05, and included the number of ATAC fragments as a latent variable. We also filtered context-relevant motifs using the regulator expression levels. The regulator associated with the motif needed to be expressed in at least 10% of the cells in the cluster.

### Trajectory Inference Analysis

Pseudotime trajectory analysis was conducted using Monocle v3^51,52^. Briefly, RNA counts were preprocessed using the preprocess_cds function with the PCA method. To correct batch effects, we used the align_cds function^53^ with the sample as an alignment group and a residual formula including the sequencing batch, brain region, and mitochondrial genes percentage as covariates. Nuclei were then projected onto the UMAP space^54^ and divided into clusters and partitions with the cluster_cells function^55^. Subsequently, an unsupervised trajectory was identified using the SimplePPT (Probabilistic Pseudotime Trajectory) algorithm implemented in the learn_graph function^56^, with the root set according to the expression of immature cellular markers. Pseudotime values were automatically assigned to individual cells based on their distance along the trajectory relative to the root node. Genes with differential expression along the trajectory were calculated using the graph_test function, with adjusted P-value less than 0.05 and arranged according to Moran’s I statistic. For each of these genes, differential expression between tissue conditions was assessed using the fit_models and coefficient_table functions, using a regression test with an adjusted P value less than 0.05, minimum fold-change of 1.5, and with the number of counts, sequencing batch, brain region and mitochondrial gene percentages as covariates.

### Inference of regulatory links

We used a regression approach based on random forests to find open chromatin regions associated with gene regulation. The goal was to predict the expression levels of the target gene based on the accessibility of ATAC peaks (i.e. cis-regulatory elements) around the gene promoter. Considering that a target gene *g* has *n* chromatin peaks, the linear regression was defined as:

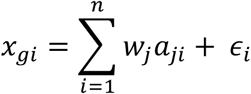

where *x_gi_* is the expression of *g* in cell *i*, *a_ji_* is the accessibility of a given peak *j* in cell *i*, *w_j_* is the weight and *ε_i_* is the noise term. For the regression, we tested all peaks located in a 500kb window around the gene’s transcriptional start site (TSS), to include both proximal and distal regulatory elements. The optimal weights, which were associated with the importance of the cis-regulatory regions as predictors of gene expression, were determined using random forests. Specifically, the regression was implemented using the GENIE3 R package^49^, setting the number of trees to 200, and peaks with weights in the top 5% (95% percentile) for each gene were selected. We used the Encode annotation^57^ downloaded from https://screen.encodeproject.org to identify open chromatin regions with potential enhancer activity. Encode enhancers are classified based on their relative location to the TSS as proximal enhancer-like sequences (pELS) if located within 200bp < TSS ≤ 2kb, or distal enhancer-like sequence (dELS) when located more than 2kb from a TSS. We used both pELS and dELS datasets for annotation.

## Acknowledgments

We thank the patients and their families who participated in this study. We thank Claudia Kleimann, Steven Hebert, Bhavyaa Chandarana and Tomas V. Waichman for advice on single-cell analysis; Damien Faury for sharing the nuclei extraction protocol; Danielle C. Bruno for discussions regarding the tissue dissociation. This work was supported by the following grants: Fundação de Amparo à Pesquisa do Estado de São Paulo (FAPESP) Young Investigator grant 2019/07382-2 (D.F.T.V); the Chan Zuckerberg Initiative DAF, an advised fund of the Silicon Valley Community Foundation grant DAF2021-237598 (D.F.T.V.); FAPESP doctoral fellowship 2022/01530-2 (I.C.G); Coordenação de Aperfeiçoamento de Pessoal de Nível Superior (CAPES) master fellowship (L.A.M.); Conselho Nacional de Pesquisa (CNPq) grant 311923/2019-4 (I.L.-C.); FAPESP doctoral fellowship 2020/06168-4 (J.C.G.), FAPESP regular grant 2019/08259-0 (F.R.), and FAPESP Research Innovation and Dissemination Center grant 2013/07559-3 (F.C.).

## Author contributions

I.C.G. and D.F.T.V. designed the study. L.A.M. and M.L. performed single-cell assays. I.C.G. and D.F.T.V. performed data analyses. P. A., J.C.G., C.L.Y., E.G., H.T., M.K.M.A., J.C.G, F.C., and I.L-C contributed to clinical sample collection. F.R. performed histopathological analyses. I.C.G. and D.F.T.V. wrote the manuscript. All authors reviewed and approved the final version of the manuscript.

